# vqtl: An R package for Mean-Variance QTL Mapping

**DOI:** 10.1101/149377

**Authors:** Robert W. Corty, William Valdar

## Abstract

We present `vqtl`, an `R` package for mean-variance QTL mapping. This QTL mapping approach tests for genetic loci that influence the mean of the phenotype, termed mean QTL, the variance of the phenotype, termed variance QTL, or some combination of the two, termed mean-variance QTL. It is unique in its ability to correct for variance heterogeneity arising not only from the QTL itself but also from nuisance factors, such as sex, batch, or housing. This package provides functions to conduct genome scans, run permutations to assess the statistical significance, and make informative plots to communicate results. Because it is inter-operable with the popular `qtl` package and uses many of the same data structures and input patterns, it will be straightforward for geneticists to analyze future experiments with `vqtl` as well as re-analyze past experiments, possibly discovering new QTL.

## INTRODUCTION

Traditional quantitative trait locus (QTL) analyses have focused on discovering “mean QTL” (mQTL), regions of the genome where allelic variation drives heterogeneity of phenotype mean, while assuming that the residual variance, that is, the intrinsic stability or noisiness of the phenotype, is identical for every individual in the mapping population. It has long been recognized, however, that the residual variance is itself heritable (Falconer 1965; Lynch and Walsh 1998), a phenomenon that has been described theoretically (Hill and Zhang 2004; Hill and Mulder 2010), demonstrated in inbred model organisms (Sorensen *et al.* 2015) and crops (Yang *et al.* 2012b), and exploited in livestock improvement efforts (Mulder *et al.* 2008; Ibáñez-Escriche *et al.* 2008). Correspondingly, several groups have proposed statistical methods for mapping QTL controlling the extent of this residual variance, these sometimes termed “variance QTL” (vQTL) (Paré *et al.* 2010; Rönnegård and Valdar 2011, 2012; Cao *et al.* 2014; Soave and Sun 2017; Dumitrascu *et al.* 2018). However, although detection of vQTL has started to enter the mainstream of genetic analysis (Yang *et al.* 2012a; Hulse and Cai 2013; Ayroles *et al.* 2015; Forsberg *et al.* 2015; Wei *et al.* 2016; Wang and Payseur 2017; Wei *et al.* 2017), statistical tools for this purpose remain heterogeneous.

We have developed a standardized method for QTL mapping in experimental crosses, in particular F2 intercrosses and backcrosses, that simultaneously models mean and variance effects in order to detect mQTL, vQTL and a generalization of the two that we term “mvQTL”. Our approach, which we term “mean-variance QTL mapping”, is based on a double generalized linear model (DGLM) (Smyth 1989), following the proposed use in this context by Rönnegård and Valdar (2011). In the first of two companion articles, we characterize the method and competitors in the setting where variance heterogeneity is driven by a background factor, such as sex, batch or housing, and show that modeling these (external) variance effects improves power to detect mQTL, vQTL and mvQTL (Corty and Valdar 2018+). In the second companion paper, we demonstrate the approach on two existing datasets and discover new mQTL and vQTL (Corty *et al.* 2018+).

Here, we provide a practical guide to the approach using its associated R package vqtl, which is currently suitable for F2 intercrosses and backcrosses, and is inter-operable with the well-established mean QTL-oriented package for this purpose, qtl (Broman *et al.* 2003). First, to generate illustrative data, we simulate an F2 intercross and four phenotypes: one phenotype determined entirely by random noise, and one with each of the three kinds of QTL. On each phenotype we then conduct a genome scan using standard approximations to interval mapping (Lander and Botstein 1989; Martínez and Curnow 1992), and mean-variance QTL mapping, which includes a test for mQTL, a test for vQTL, and a test for mvQTL. The association statistics of all four tests are initially plotted in LOD score units, with drawbacks of this plotting unit discussed; then permutation scans are used to determine empirically-adjusted *p*-values, and plotting in these units is shown to to make the results of the four tests more easily comparable. Plots are then described that communicate effects that led to the QTL’s detection, and the bootstrap is used to estimate its confidence interval. Last, we benchmark performance, using one of the datasets examined in Corty *et al.* (2018+) to report how computation time varies with marker density and number of permutations.

## EXAMPLE DATA: SIMULATED F2 INTERCROSS

To illustrate the use of the `vqtl` package, we first simulated an example F2 intercross using the `R` package qtl (Broman *et al.* 2003), on which vqtl is based. This cross consisted of 200 male and 200 female F2 offspring, with 3 chromosomes of length 100 cM, each tagged by 11 equally-spaced markers and estimated genotype probabilities at 2cM intervals with `qtl`’s hidden Markov model. We then generated four phenotypes:

1. `phenotype1` consists only of random noise and will serve as an example of negative results for all tests.
2. `phenotype2` has an mQTL that explains 4% of phenotype variance at the center of chromosome one.
3. `phenotype3` has a vQTL at the center of chromosome two. This vQTL acts additively on the log standard deviation scale, and results in residual standard deviation of [0.8, 1, 1.25] for the three genotype groups.
4. `phenotype4` has an mvQTL at the center of chromosome three. This mvQTL has a mean effect that explains 2.7% of phenotype variance and a variance effect that acts additively on the standard deviation scale, resulting in residual standard deviation of [0.85, 1, 1.17] for the three genotype groups.

We additionally consider `phenotype1x` through `phenotype4x`, which have the same type of genetic effects as `phenotype1` through `phenotype4`, but have the additional feature that females have greater residual variance than males. All the same analyses and plots that are shown for `phenotype1` through `phenotype4` are shown for `phenotype1x` through `phenotype4x` in the appendix.

## SCAN THE GENOME

The central function for genetic mapping in package `qtl` is `scanone` (Broman *et al.* 2003). Analogously, the central function for mean-variance QTL mapping in package `vqtl` is `scanonevar`, building on an early version of `scanonevar` in package qtl. It takes three required inputs:

1. `cross` is an object that contains the genetic and phenotypic information from an experimental cross, as defined in package `qtl`.
2. `mean.formula` is a two-sided formula, specifying the phenotype to be mapped, the covariates to be corrected for, and the QTL terms to be fitted, with keywords `mean.QTL.add` and `mean.QTL.dom`
3. `var.formula` is a one-sided formula, specifying the variance covariates to be corrected for as well as the QTL terms to be fitted, using keywords `var.QTL.add` and `var.QTL.dom`.

For example, to scan a phenotype named p1, we run:

`scanonevar(`

`cross = test_cross,`

mean.formula = p1 ^~^ sex + mean.QTL.add + mean.QTL.dom, var.formula = ^~^ sex + var.QTL.add + var.QTL.dom

)

At each locus in turn, this function tests for the presence of an mQTL, a vQTL, and an mvQTL. The basis of these tests is a comparison between the fit of an alternative model of the form

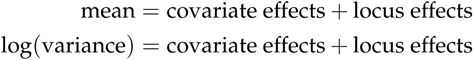

with a null model that omits specific terms: for the mQTL test, the null model omits locus effects on phenotype mean; for the vQTL test, the null omits the locus effects on phenotype variance; and for the mvQTL test, the null omits locus effects on both mean and variance. (Note that the mQTL test in mean-variance QTL mapping is different from the traditional test: the traditional test does not have variance predictors of any kind in either null or alternative models.)

### LOD scores and nominal p-values

Each type of test (mQTL, vQTL, and mvQTL) yields two association statistics: the LOD score, and the (nominal) *p*-value. The LOD is a raw measure of association equal to the base 10 logarithm of the likelihood ratio (LR) between the fitted alternative and null models. Higher values indicate greater association when considered across loci for the same type of test; but LOD scores between different types of tests, namely between mvQTL test vs either mQTL or vQTL tests, are not readily comparable. The *p*-value, which is comparable between different types of tests, transforms the LOD score to take account of the number of parameters being fit: it is calculated from the asymptotic distribution of 2 log_*e*_ (LR) under the null model, namely the *χ*2 distribution with degrees of freedom equal to the difference in the number of parameters between the alternative and null models.

The *p*-values described above, however, are nominal: they do not take into account multiple testing across the genome. They also rely on asymptotic theory that assumes the underlying phenotype being residually normal; this may not always be the case and when violated will lead to inflated significance. More robust *p*-values that are corrected for genomewide significance via control of the family-wise error rate (FWER) can be obtained empirically, through a permutation procedure described below.

### Robust, genomewide-adjusted p-values

To calculate the empirical, FWER-controlled *p*-value of each test at each locus we advocate use of a permutation procedure (Corty and Valdar 2018+). Like previous work on permutation-based thresholds for genetic mapping (Churchill and Doerge 1994; Carlborg and Andersson 2002), this procedure sidesteps the need to explicitly estimate the effective number of tests.

In brief, this approach involves conducting many genomes scans on pseudo-null data generated through permutation to maintain as much of the character of the data as possible, while breaking the tested phenotype-genotype association. Specifically, the design matrix of the QTL is permuted in the mean portion of the mQTL alternative model, the variance portion of the vQTL alternative model, and in both portions of the mvQTL alternative model.

For each test (mQTL, vQTL, and mvQTL), the highest observed test statistic is extracted from each permutation scan and the collection of statistics that results is used to fit a generalized extreme value (GEV) density (Stephenson 2002; Dudbridge and Koeleman 2004; Valdar *et al.* 2006). The observed LOD scores from the genome scan are then transformed by the cumulative distribution function of the extreme value density to estimate the FWER-controlling *p*-values. This approach is implemented in the function, `scanonevar.perm`, which requires two inputs:

1. `sov` is the `scanonevar` object, the statistical significance of which will be assessed through permutation.
2. `n.perms` is the number of permutations to conduct.

The object returned by `scanonevar.perm` is a `scanonevar` object with two additional pieces of information: an empirical *p*-value for each test at each locus and the per-permutation maxima that were used to calculate those *p*-values. These FWER-corrected *p*-values are straightforwardly interpretable: *p* = 0.05 for a specific test at a specific locus implies that in 5% of similar experiments where there is no true genotype-phenotype association, we would expect to observe some locus with this much or more evidence of association in this test.

Accurate estimation of the FWER-controlled *p*-values requires many permutation scans: traditionally recommended is 1,000 (*e.g.*, Churchill and Doerge 1994; Carlborg and Andersson 2002), although the efficiency gain of using the GEV rather than raw quantiles means that fewer may be adequate in practice (Valdar *et al.* 2006). These permutation scans can be run on multiple processors by specifying the optional `n.cores` argument, which defaults to the total number of cores on the computer minus 2. On an Intel Core i5, running 100 permutations on this dataset takes about five minutes. When many phenotypes are studied, or if faster runtimes are needed, these permutation scans can be broken into groups with different values for `random.seed`, run on separate computers, and combined with the c function. This function combines the permutations from all the inputted scans, re-estimates the extreme value density, re-evaluates the observed LOD scores in the context of new extreme value density, and returns a new `scanonevar` object with more precisely estimated empirical *p*-values.

### Reporting and plotting genome scans

The results of scanonevar can be plotted by calling plot on the scanonevar output object. This produces a publication-quality figure that shows the association of the phenotype for each location in the genome as different colors for type of test, with y-axis scale being specified by the user, via option `plotting.units` as the LOD (Figure 1), nominal *p*-value, or, provided permutations have been run, empirical, FWER-controlling *p*-value (Figure 2). Of the available y-axis scales, we recommend using the FWER-controlled *p*-values since this scale puts all tests on a level-footing (unlike the LOD), and allows direct identification of genomewide significance and thereby relevance (unlike the nominal *p*-value).

**Figure 1.**
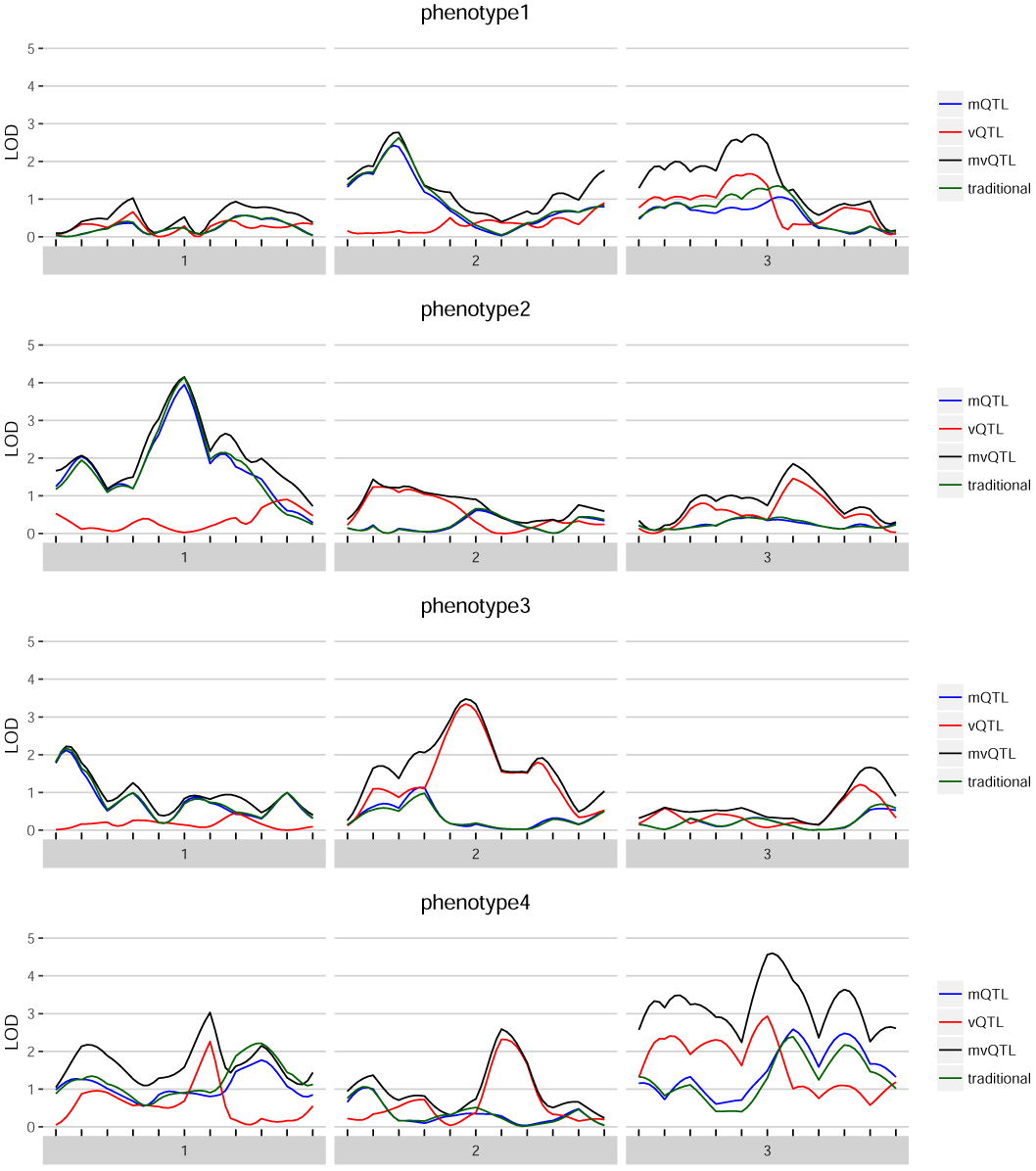
For each of the four simulated phenotypes, the genome scan shows the LOD score of each test — mQTL, vQTL, and mvQTL — in blue, red, and black, respectively. The traditional test is in green and globally similar to the mQTL test.

**Figure 2.**
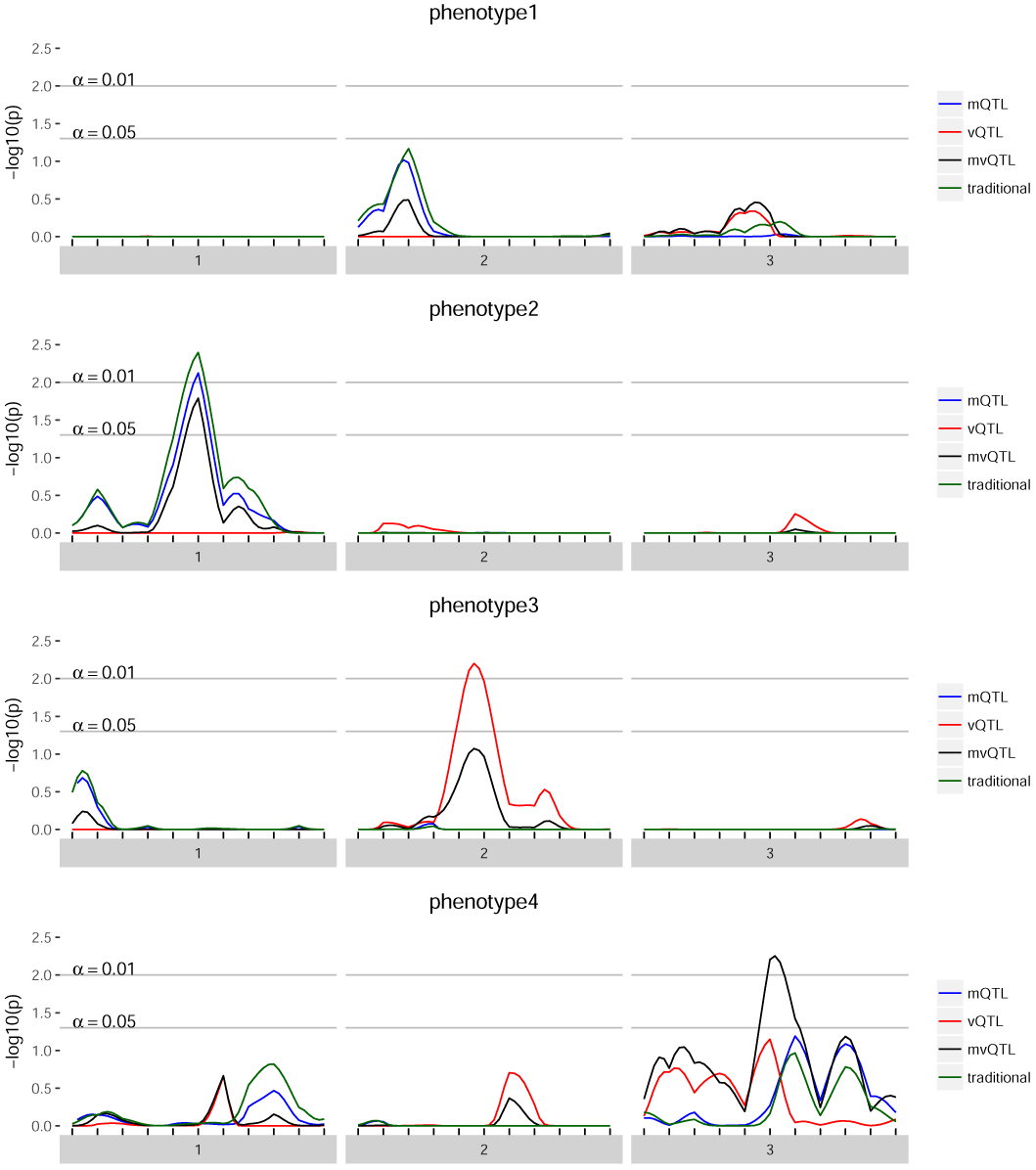
For each of the four simulated phenotypes, the genome scan shows the -log10 of the FWER-corrected *p*-value for each test — mQTL, vQTL, and mvQTL — in blue, red, and black, respectively. The traditional test is in green and globally similar to the mQTL test. A value of 2 implies that the quantity of evidence against the null is such that we expect to see this much or more evidence once per hundred phenotypes no QTL.

Calling `summary` on the output of `scanonevar` produces a summary of how the scan was conducted and what the results were.

## COMMUNICATE SIGNIFICANT FINDINGS

Having identified interesting QTL, we want to visualize the their estimated genetic and covariate effects. Because the vqtl package models effects for both mean and variance, existing plotting utilities are not able to display the entirety of the modeling results. To understand and communicate the results of a vqtl scan at one particular locus, we developed the `mean_var_plot`. This plot illustrates how the mean sub-model and variance sub-model of the DGLM fit the data at a given locus.

In each `mean_var_plot` in Figure 3, the location of the dot shows the estimated mean and standard deviation of each genotype group, with the mean indicated by the horizontal position and the standard deviation indicated by the vertical position. The horizontal lines extending to the left and right from each dot show the standard error of the mean estimate, and the vertical lines extending up and down from each dot show the standard error of the standard deviation estimate. There are two types of grouping factors considered by the function `mean_var_plot_model_based`: (1) `focal.groups` are groups that are modeled and the prediction for each group is plotted. For example, a genetic marker is the `focal.group` in each plot in Figure 3; `D1M1` in the top left, `D1M6` in the top right, `D2M6` in the bottom left, and `D3M6` in the bottom right. (2) `nuisance.groups` are groups that are modeled, but then averaged over before plotting. When there are many grouping factors thought to play a role in determining the mean and variance of an individual’s phenotype, such as sex, treatment, and batch, we recommend putting just one or two in `focal.groups` and the others in `nuisance.groups` for clarity, cycling through which are displayed to gain a thorough understanding of the factors that influence the phenotype.

**Figure 3.**
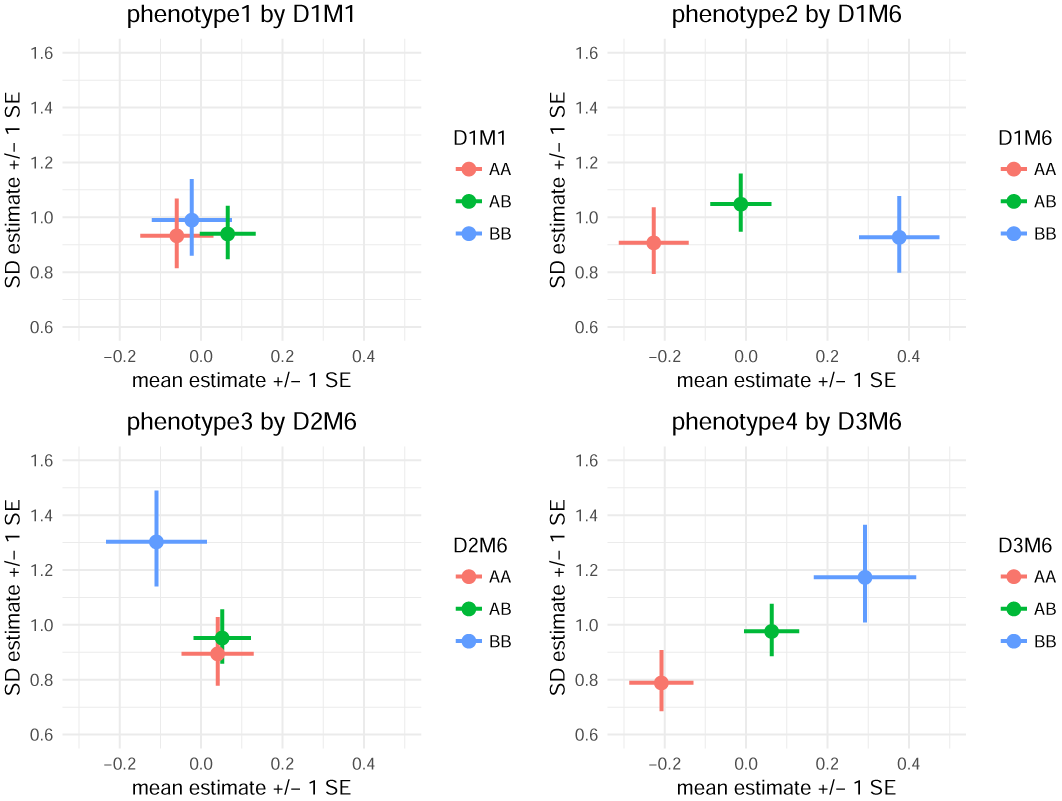
`mean_var_plots` show the estimated genotype effects at a locus with mean effects on the horizontal axis and variance effects on the vertical axis. Horizontal lines indicate standard errors for mean effects and vertical lines indicate standard errors for variance effects. For `phenotype1`, the pattern of overlapping estimates and standard errors is consistent with the fact that there are no genetic effects, and the *p*-value was not statistically significant at any locus. For `phenotype2`, the pattern of horizontal, but not vertical, separation visually illustrates the identified mQTL. For `phenotype3`, the pattern of vertical, but not horizontal, separation visually illustrates the identified vQTL. For `phenotype4`, the pattern of two-dimensional separation illustrates an mvQTL.

Additional plotting utilities, `phenotype_plot`, `effects_plot` and `mean_var_plot_model_free` are described in the online documentation, available on CRAN.

## ESTABLISH A CONFIDENCE INTERVAL FOR THE QTL

Last, to assess the genetic precision of a discovered QTL for bioinformatic follow-up, the function `scanonevar.boot` estimates confidence intervals via the non-parametric bootstrap (Visscher *et al.* 1996). This function takes, as arguments, a `scanonevar` object, the type of QTL detected, the name of the chromosome containing the QTL, and `num.resamples`, the number of bootstrap resamplings desired. As with `scanonevar.perm`, the `n.cores` argument can be used to spread the bootstraps over many computational cores and defaults to the number of cores available minus two, and bootstraps can be run on separate computers and combined with c to increase the precision of the estimate of the confidence interval.

We recommend 1000 resamples to establish 80% and 90% confidence intervals. With the datasets simulated here, it takes 20 minutes to run 1000 bootstrap resamples on an Intel core i5.

## PERFORMANCE BENCHMARKS

By far, the most computationally-intensive step in the mean-variance QTL mapping process is the assessment of genome-wide statistical significance by permutation. The original genome scan is much faster, because it involves only a single scan, and the bootstrap is much faster because it involves only a single chromosome.

For the first benchmark, we ran `scanonevar.perm` on the data from Kumar *et al.* (2013) and Corty *et al.* (2018+), which contains 244 individuals and 582 loci, varying the number of permutations desired and the number of computer cores used. For a given number of cores, the relationship between time and the number of permutations is linear (Figure 4), the slope depending on the number of cores and ranging from ≈6.3 seconds per permutation with 4 cores to ≈1.2 second per permutation with 32 cores.

**Figure 4.**
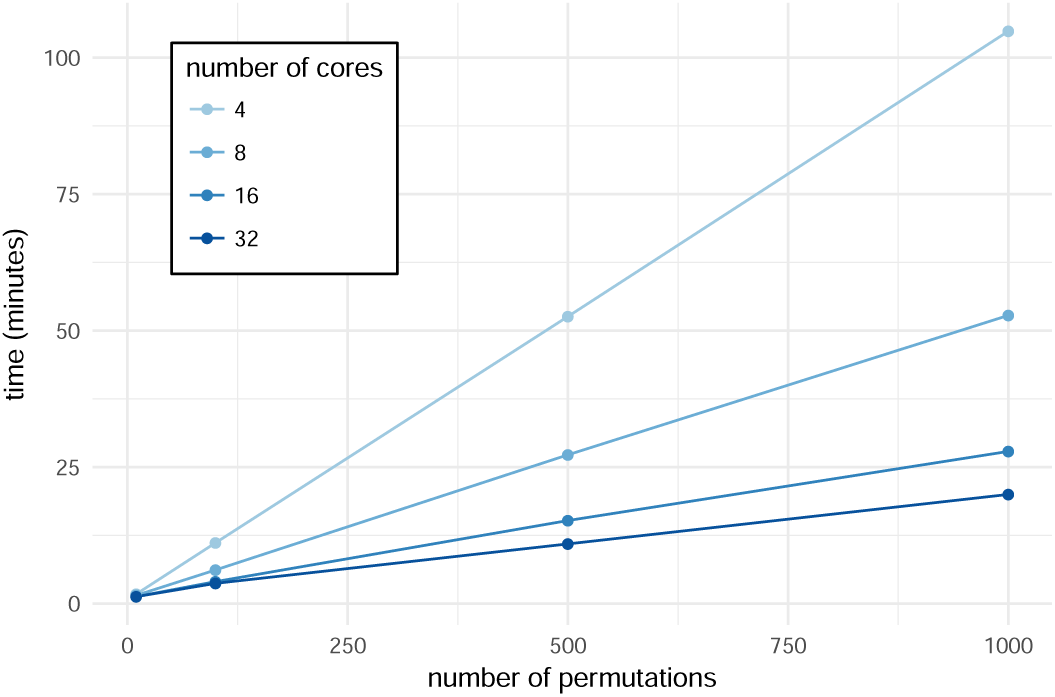
Time taken to run `scanonevar.perm` on the data from Kumar *et al.* (2013) which contains 244 individuals and 582 loci, varying the number of permutations desired and the number of computer cores used. For a given number of cores, there is a linear relationship between number of permutations conducted and time required. The slope the the line indicates time required per permutation and is dependent on the number of cores, ranging from ≈6.3 seconds per permutation with 4 cores to ≈1.2 second per permutation with 32 cores.

For the second benchmark, we ran `scanonevar.perm` on simulated data, always conducting 1000 permutations and using 32 cores, but varying the number of individuals in the mapping population and the number of markers in the genome. For a given population size, there is a slightly curvilinear relationship between number of markers and time required (Figure 5), which reflects a linear increase in the time taken to conduct the permuted genome scans plus an increase in the time taken for “bookkeeping” tasks like organizing and reshaping genetic data. The slope (minutes per locus) depends on the population size, ranging from ≈1.4 seconds per locus with a population of size 100 to ≈3.3 seconds per locus with a population of size 800.

**Figure 5.**
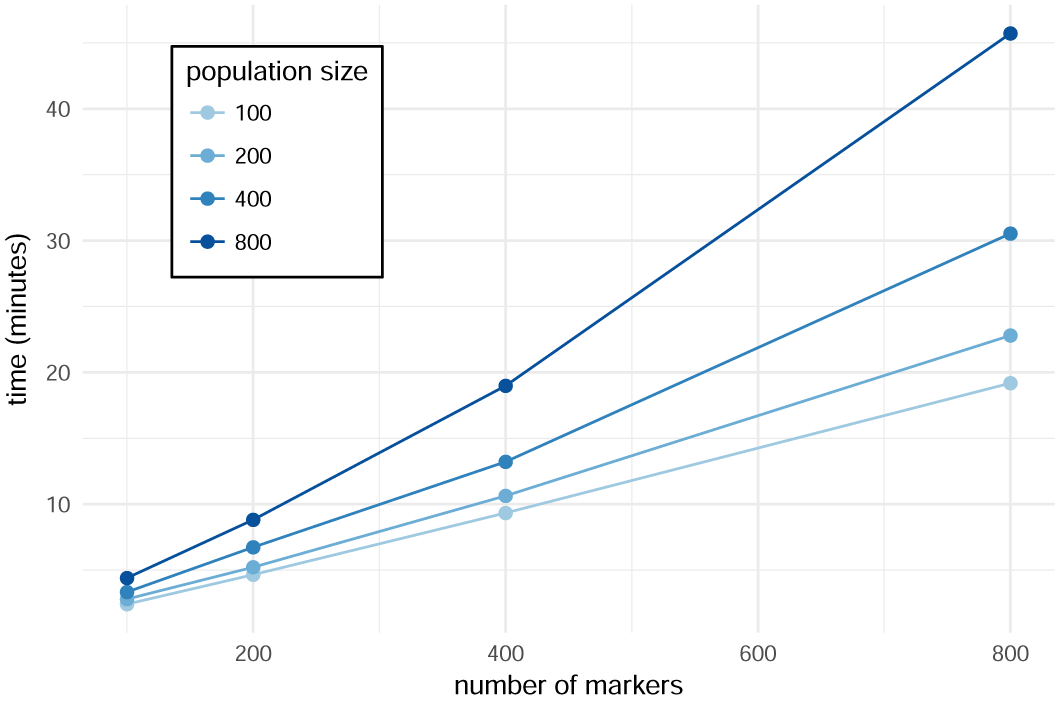
Time taken to run 1000 permutation scans on 32 cores on simulated data using `scanonevar.perm`, varying the number of individuals in the mapping population and the number of markers in the genome. For a given population size, there is a slightly supra-linear relationship between number of markers and time required. The average slope of the line indicates the average time required per locus and is dependent on the population size, ranging from ≈1.4 seconds per locus with a population of size 100 to ≈3.3 seconds per locus with a population of size 800.

Based on these benchmarks, the workflow presented here is practical for QTL mapping F2 intercross and similar populations on modern, multi-core scientific computers. Populations with many recombinations, where dense genotyping arrays that interrogate *>* 10,000 loci, could not be practically analyzed with package `vqtl` in this way, although it is likely that statistical and computational steps could be taken to make such studies feasible: statistically, techniques could be used that allow for large-scale analysis without permutation testing (Efron 2004); computationally, the software could be modified to run on a computer cluster, rather than on a single computer (Jette and Grondona 2003; Marchand 2017).

## CONCLUSION

We have demonstrated typical usage of the `R` package `vqtl` for mean-variance QTL mapping in an F2 intercross. This package is appropriate for crosses and phenotypes where genetic factors or covariates or are known or suspected to influence phenotype variance. In the case of genetic factors, they can be mapped, as illustrated in one companion article (Corty *et al.* 2018+). In the case of covariates, they can be accommodated, which can increase power and improve false positive rate control, as illustrated in another companion article ((Corty and Valdar 2018+)).

## RESOURCES

The scripts used to simulate genotypes and phenotypes, conduct the genome scans, and plot the results are available as a public, static Zenodo repository at DOI:10.5281/zenodo.1336302. The package `vqtl` and its documentation are freely available on `CRAN` at https://CRAN.R-project.org/package=vqtl.

## ACKNOWLEDGMENTS

This work was funded by National Institutes of General Medical Sciences grants R01-GM104125 (RWC,WV), R35-GM127000 (RWC,WV), T32-GM067553 (RWC); National Heart, Lung and Blood Institute grant R21 HL126045 (RWC,WV); National Library of Medicine grant T32-LM012420 (RWC); and a National Institute of Mental Health grant F30-MH108265 (RWC).

**Figure A1.**
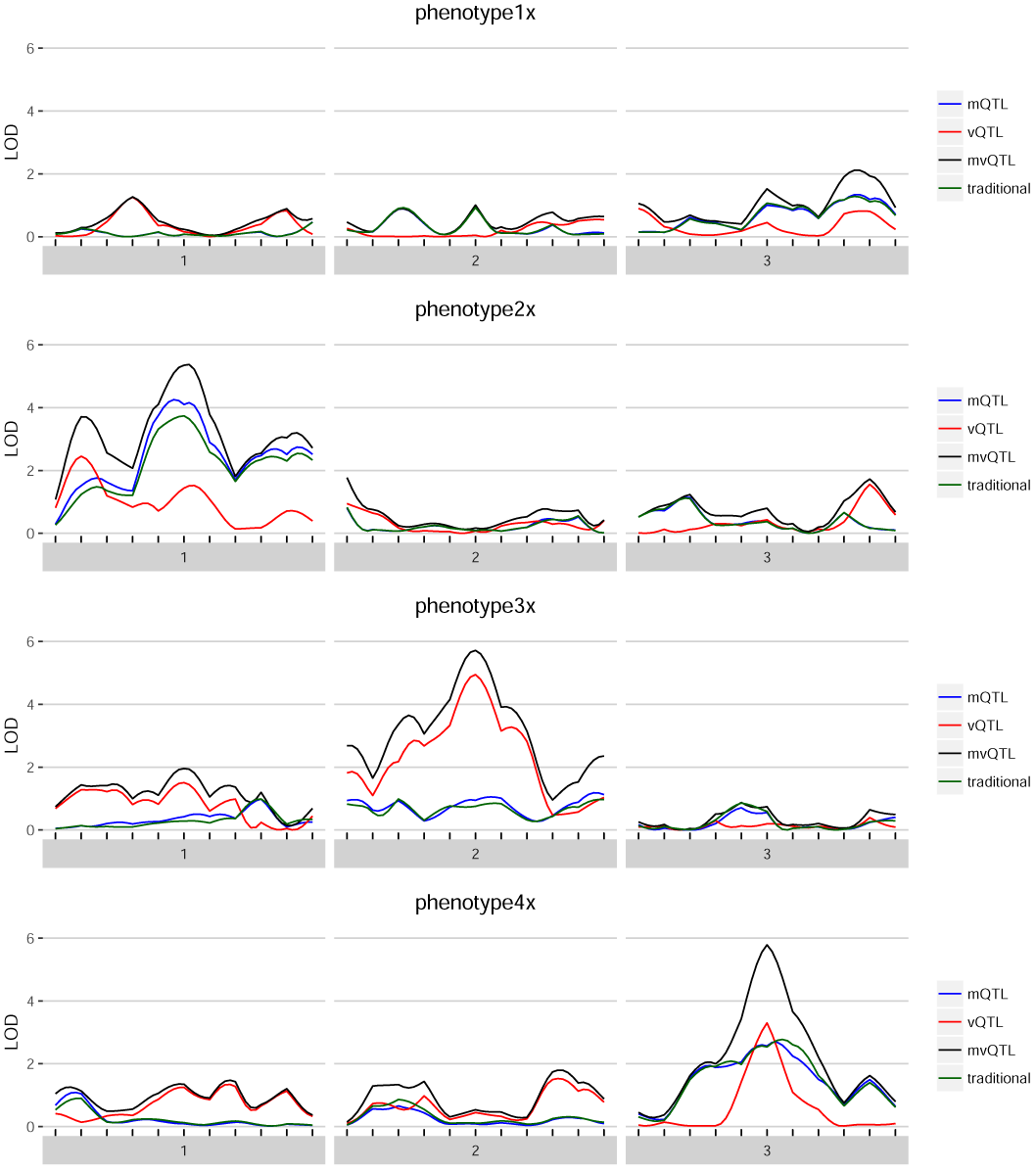
For each of the four simulated phenotypes with background variance heterogeneity, the genome scan shows the LOD score of each test – mean, variance, and joint – in blue, red, and black, respectively. The traditional test is in green and globally similar to the mean test.

**Figure A2.**
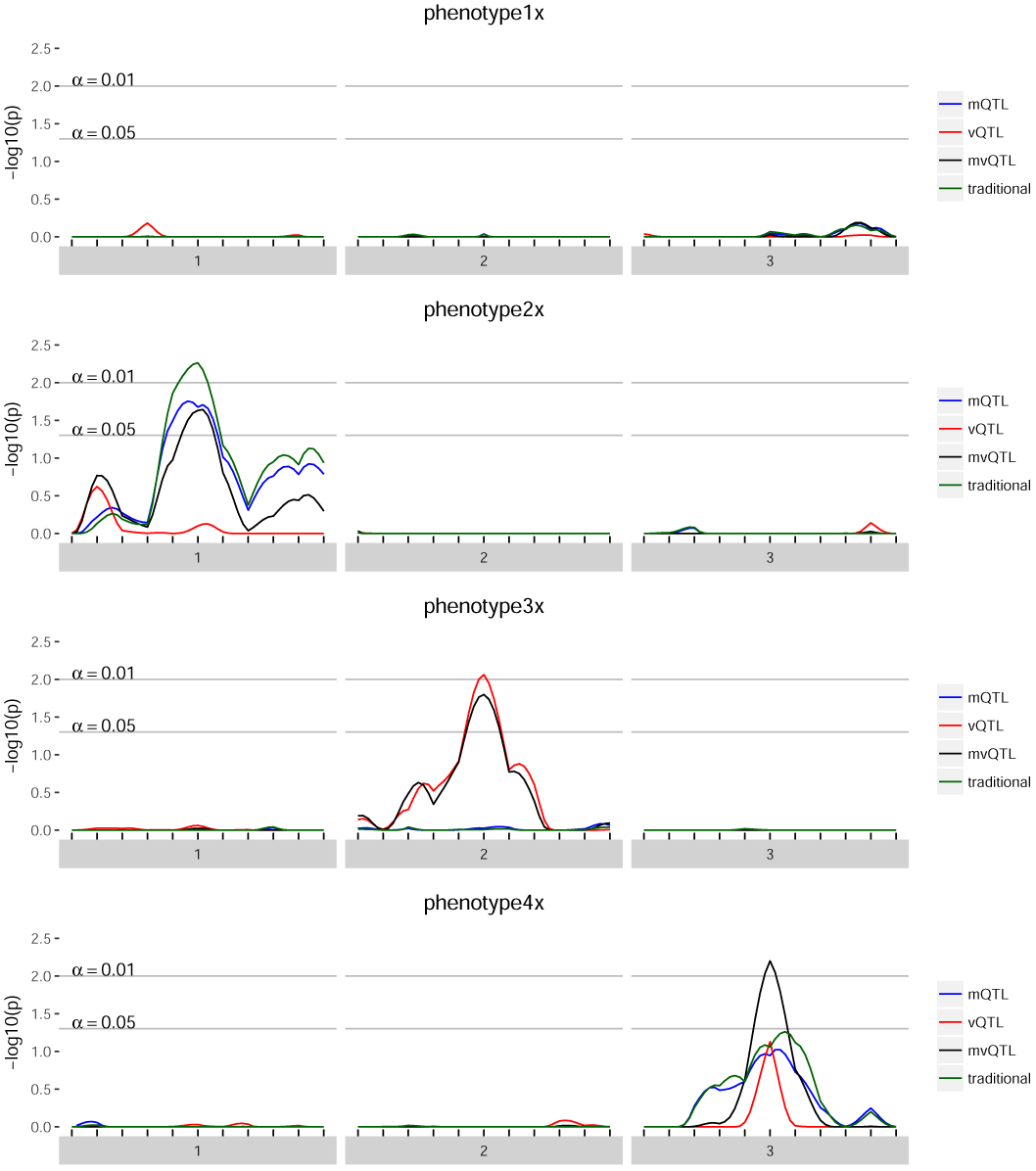
For each of the four simulated phenotypes with background variance heterogeneity, the genome scan shows the - log10 of the FWER-corrected *p*-value of each test – mean, variance, and joint – in blue, red, and black, respectively. Thus, a value of 3 implies that the quantity of evidence against the null is such that we expect to see this much or more evidence once per thousand genome scans when there is no true effect.

**Figure A3.**
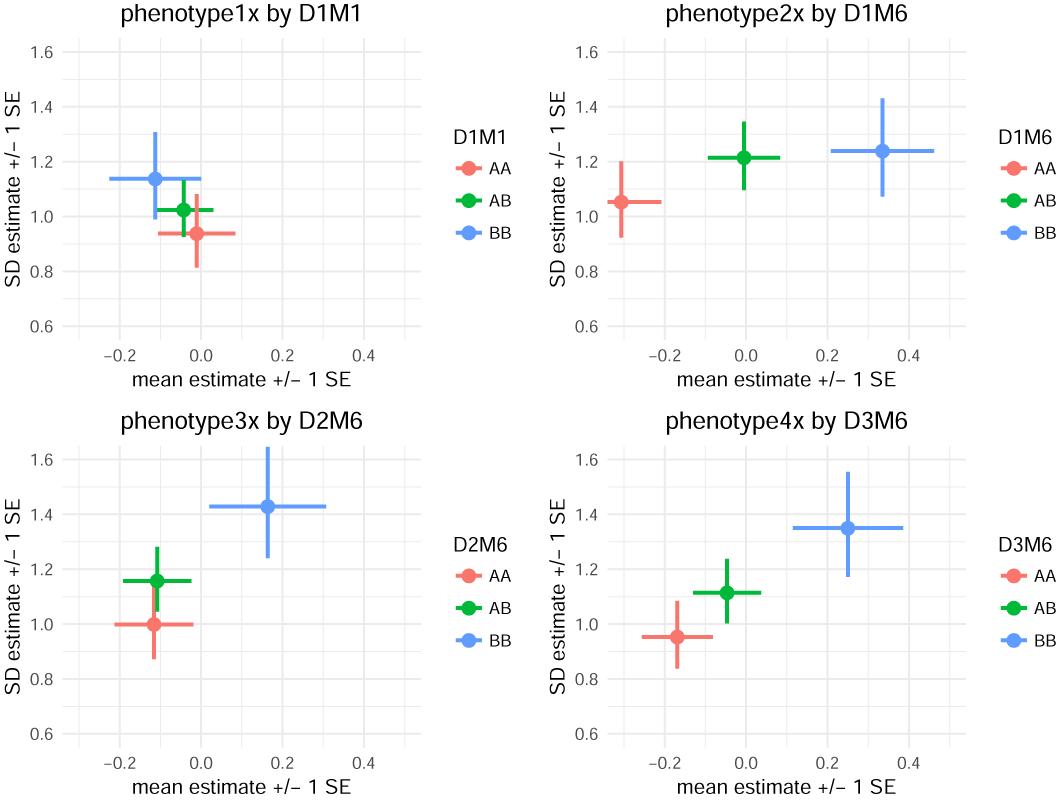
`mean_var_plots` show the estimated genotype effects at a locus, with mean effects on the horizontal axis and variance effects on the vertical axis. Horizontal lines indicate standard errors for mean effects and vertical lines indicate standard errors for variance effects. For `phenotype1x`, the pattern of overlapping estimates and standard errors is consistent with the fact that there are no genetic effects, and the *p*-value was not statistically significant at any locus. For `phenotype2x`, the pattern of horizontal, but not vertical, separation visually illustrates the identified mQTL on a background of variance heterogeneity. For `phenotype3x`, the pattern of vertical, but not horizontal, separation visually illustrates the identified vQTL on a background of variance heterogeneity. For `phenotype4x`, the pattern of two dimensional separation without either total horizontal or vertical separation illustrates an mvQTL with neither mean nor variance effect strong enough to define an mQTL or vQTL on a background of variance heterogeneity.

